# Sensitivity of single units in the LSO of decerebrate cat to sinusoidally amplitude modulated tones

**DOI:** 10.1101/515270

**Authors:** Nathaniel T. Greene, Kevin A. Davis

**Author notes:** Send Correspondence to: Nathaniel T Greene, Ph.D., University of Colorado School of Medicine, Department of Otolaryngology, Research Complex 1-N, Rm 7104, 12631 E. 17^th^ Ave, MS B205, Aurora CO 80045, (work).

## Abstract

Fluctuations in amplitude are a common component of behaviorally important sound stimuli. Amplitude modulation (AM) is encoded by the peripheral auditory system in the timing of discharge spikes, and, more centrally, in the discharge rate. The mechanism producing this transformation from a time- to rate-based code is not known, but recent modeling efforts have suggested a role for neurons with response characteristics consistent with cells in the lateral superior olive (LSO). The responses of single units in the LSO of unanesthetized decerebrate cat were recorded to monaural sinusoidally amplitude modulated (SAM) tones by systematically varying sound level and modulation frequency (f_m_), and are described in terms of synchronization to the envelope and average discharge rate as a function of f_m_. LSO units typically synchronize strongly to low f_m_, and discharge preferentially (i.e. more strongly) over a small range of f_m_ in response to low level SAM tones. At higher sound levels synchronization decreases and response rate increases until most or all modulation in the response is lost. These results contrast with responses recorded in the barbiturate-anesthetized cat, which tend to respond to most low-frequency modulations, and are consistent with LSO as an intermediate processing stage between the peripheral temporal- and central rate-based code for AM sounds.

## INTRODUCTION

Variations in the amplitude of sounds are found in many commonly encountered sounds (Singh and Theunissen 2003), and are utilized for several functions performed by the auditory system including the perception of pitch (Plack 2005), analysis of vocalizations (Rosen 1992; Steeneken and Houtgast 1980), auditory scene analysis (Bregman et al. 1990), and localization of high frequency sounds (Henning 1974a; b). Physiological studies suggest that amplitude modulated (AM) stimuli are represented in the timing of discharge spikes in the auditory periphery, while more centrally, some units respond selectively to particular modulation frequencies (f_m_) (for review, Frisina 2001; Joris et al. 2004). The precise mechanism effecting this transformation from a time-to rate-based code is not known.

Four competing models of AM sensitivity in the auditory system have been developed to explain rate-based tuning in the auditory midbrain (for review: Davis et al. 2010). In general, these models can be segregated based upon the neural population in cochlear nucleus (CN) through which information is transmitted from auditory nerve (AN) to inferior colliculus central nucleus (ICC). Most (3/4) are initiated by CN chopper cells (Dicke et al. 2007; Hewitt and Meddis 1994; Langner 1981), which respond to stimuli with highly regular and reliable discharges (Young et al. 1988). The fourth model (Nelson and Carney 2004) does not require the regular discharges of chopper cells, and instead uses parameters from bushy cells, which produce primary-like (i.e. AN-like) rather than chopper discharge patterns (Rhode et al. 1983). The results of this latter model suggest that same-frequency inhibitory and excitatory (SFIE) inputs, which are ubiquitous within the auditory system (e.g. Caspary et al. 1994; Ferragamo et al. 1998), can produce a discharge rate response tuned to a particular range of f_m_ without requiring specialized discharge characteristics. Bushy cell outputs are relayed to the ICC through cells in the superior olivary complex (SOC) and ventral nucleus of the lateral lemniscus (VNLL) to ICC (Cant and Benson 2003; Cant and Casseday 1986), suggesting that SOC cells could contribute to AM processing. Responses of VNLL cells to AM stimuli, which provides a major source of inhibitory input to ICC (Saint Marie et al. 1997; Whitley and Henkel 1984; Winer et al. 1995), have received considerable attention recently, which has revealed tuned rate functions (Batra 2006; Zhang and Kelly 2006). In contrast, investigations of the responses of SOC cells to AM stimuli have been more limited.

Cells in the SOC are among the first cells to receive information from both ears. In particular, cells in the lateral superior olive (LSO) receive excitatory input from ipsilateral spherical bushy cells, and inhibitory input from contralateral globular bushy cells, via the ipsilateral medial nucleus of the trapezoid body (MNTB) (Glendenning et al. 1985; Glendenning and Masterton 1983; Tolbert et al. 1982), thus are sensitive to interaural level differences (ILDs; e.g. Boudreau and Tsuchitani 1968; Tollin et al. 2008). LSO cells then send excitatory projections to contralateral, and inhibitory projections to ipsilateral ICC (Glendenning et al. 1992; Glendenning and Masterton 1983). In addition to this classic bilateral input pattern, anatomical and physiological evidence suggests an ipsilateral source of inhibition contributes to LSO responses (Caird and Klinke 1983; Greene and Davis 2012; Thompson and Schofield 2000; Wu and Kelly 1994), thus a SFIE type mechanism could produce tuned rate responses to AM stimuli in LSO cells. However, while several studies have investigated binaural processing of AM stimuli in the LSO, such as envelope interaural time difference (ITD) sensitivity (Batra et al. 1997a; b; Joris 1996; Joris and Yin 1995; 1998), few have investigated the inherent sensitivity of these cells to monaural AM stimuli.

Evidence for somewhat enhanced monaural AM rate tuning in the LSO (compared to their bushy cell inputs) was documented in the last report of the series by Joris and Yin (1998), who showed that LSO units tended to respond with higher discharge rates to low than high f_m_ SAM tones; however, units in this study were recorded under the influence of barbiturate anesthesia, which is known to affect the balance of excitation and inhibition elsewhere in the ascending auditory system (CN: Evans and Nelson 1973; ICC: Kuwada et al. 1989). Indeed, we have previously provided indirect evidence that anesthesia can affect the balance of excitation and inhibition in LSO (Greene and Davis 2012; Greene et al. 2010). Specifically, we reported that the responses of units in the unanesthetized decerebrate cat LSO to best frequency (BF) tones produce a continuous distribution of responses with four archetypes in post-stimulus time (PST) histograms: chopper-sustained (Chop-S), chopper-transient (Chop-T), primary-like (Pri-L), and onset-sustained (On-S) responses. Units displaying highly regular chopper responses predominate in the barbiturate anesthetized cat (e.g. Tsuchitani 1982), thus units recorded in unanesthetized decerebrate cat LSO shows two major differences: substantially fewer units responding with chopper-type discharge patterns are recorded, and *all* unit responses were less regular (i.e. spike discharge intervals were more variable) and reliable (i.e. spikes were not always elicited on each repetition of the stimulus presentation), consistent with recordings in the LSO of awake rabbits (Batra et al. 1997a; b). We went on to show, in a computational model of the afferent inputs to LSO, that the range of discharge patterns produced in anesthetized and unanesthetized LSO can be reproduced by varying the strength and timing of a fast ipsilaterally-driven inhibitory input to LSO that is selectively blocked by barbiturate anesthesia. This mechanism is consistent with the effects of barbiturates on glycine receptors (Lu and Xu 2002), and direct evidence of ipsilaterally evoked IPSPs in LSO, as well as their elimination following bath application of pentobarbital, has been reported in a rat slice preparation (Wu and Kelly 1994). Furthermore, limited data suggests that monaural side-band inhibition can be eliminated, and that irregular primary-like discharge patterns can be converted to regular chopper patterns through systemic application of barbiturate anesthesia in the decerebrate cat (Brownell et al. 1979). These results suggest that an ipsilaterally driven source of inhibition to LSO cells is present, consistent with a SFIE type mechanism, thus the potential exists for enhanced AM rate tuning in the unanesthetized cat LSO.

The goals of the current study were to describe the responses to monaural sinusoidally amplitude modulated (SAM) tones (with a BF tone carrier) of LSO units in decerebrate cat, and assess the ability of a sound-driven inhibitory input to reproduce this observed AM tuning via a SFIE type mechanism. Standard single-cell extracellular recording techniques were used to obtain average rate and synchronization modulation transfer functions (rMTF, sMTF) at multiple levels with respect to the unit’s best-frequency (BF) tone threshold. Unit responses are assessed as a function of their PST discharge pattern type, as both AM sensitivity (via a SFIE mechanism) and PST type (after Greene et al. 2012) may indicate the strength of inhibitory inputs onto the unit. Our results indicate that LSO units often show a prominent peak (band-pass tuning) in rMTFs, and synchronize strongly to low modulation frequencies (f_m_; low-pass tuning) at low sound levels (near the BF-tone threshold) At higher levels, response rate tends to increase at all f_m_ until the rate selectivity is lost (all-pass tuning), while synchronization decreases, sometimes decreasing preferentially at low f_m_, thus producing responses with synchronization tuned to a particular range of f_m_. Presenting AM stimuli to our previously developed model of the inputs to LSO cells, which reproduced the range of observed discharge patterns (Greene and Davis 2012), suggests that varying the strength of an ipsilaterally-driven inhibitory input can reproduce both the rate and synchronization tuning observed in decerebrate cat. All together, these data reveal that tuning at low sound levels is significantly stronger in decerebrate LSO than was reported under barbiturate anesthesia, and that LSO responses show intermediate response tuning between CN bushy cell and ICC AM responses, consistent with a multi-stage SFIE-type mechanism for transforming AM tuning from a temporal to a rate based code. These results suggest that the ascending pathway passing through the LSO may contribute to the processing of AM stimuli, likely in addition to alternative ascending pathways, such as through the VNLL, and that AM processing may be distributed across many, rather than within a single ascending pathway.

## METHODS

### Surgical Procedures

The decerebration procedure and LSO access have been described previously (Greene et al. 2010). Briefly, adult male cats (3-4kg) with infection-free ears were initially anesthetized with ketamine (40 mg/kg im) and xylazine (0.5 mg/kg im), and given atropine (0.5 mg/kg im) and dexamethasone (2 mg/kg im); supplemental doses (half the initial) were provided when indicated by the return of reflexes. In preparation for decerebration, body temperature was monitored and maintained at 39°C with a heating blanket, the cephalic vein was cannulated, and a tracheotomy was performed. Decerebration was accomplished by reflecting the skin and temporalis muscle to expose the skull, creating a fenestration over left parietal cortex, and aspirating under visual observation the underlying cortical and thalamic tissue, after which no further anesthesia was administered. The ears were laterally reflected and the external ear canals were transected to accept hollow ear bars, with which the cat’s head was secured in a stereotaxic frame. The head was rotated forward by 30°, with respect to horizontal, using a bracket secured to the forehead. Access to the left LSO was accomplished by performing a craniotomy over the midline, aspirating the underlying cerebellar tissue, and exposing the floor of the fourth ventricle and caudal extent of the cerebellar peduncle. Cats were euthanized at the end of each experiment with an overdose of sodium pentobarbital (100 mg/kg iv; Sleepaway).

### Acoustic stimuli

Sound stimuli were generated digitally with Tucker-Davis Technologies (TDT) system 3 hardware, and presented via a 16 bit digital-to-analog converter sampled at 100 kHz. The acoustic system was calibrated at the start of each experiment using a probe tube microphone (Brüel and Kjær). Most stimuli were 200 ms in duration and presented at a rate of 1 burst/s, search stimuli and detailed frequency response maps were 50 ms in duration and presented at 4 bursts/s. All stimuli were gated on/off with 10 ms rise/fall times. Tones were attenuated (TDT PA4) relative to the acoustic ceiling at that frequency and are reported in dB SPL. The spectra of noise stimuli were flat at the tympanic membrane and attenuated relative to the maximum spectrum level possible with each calibration curve (~45 dB SL). Sounds were presented using electrostatic speakers (TDT) tightly coupled to hollow ear bars.

### Recording procedure

Unit activity was recorded with platinum tipped, glass insulated platinum/iridium electrodes, using Alpha Omega hardware and software. Electrodes were advanced with a stepper-motor controlled multi-electrode positioning system (Alpha Omega EPS). The signal was amplified (10,000-30,000x) and filtered (0.3-6 kHz) using a multi-channel processor (Alpha Omega MCP). Multiple spike detecting (Alpha Omega MSD) template matching software was used to discriminate individual action potentials from the background activity. Templates for all units had error histograms with single well-defined peaks and signal-to-noise ratios greater than two-to-one (and often greater than ten to one), suggesting well-isolated single unit activity. Spike times relative to stimulus onset were recorded for on- and off-line analysis.

Electrodes were advanced into LSO using a previously described dorsal approach (Greene et al. 2010). Search stimuli, 50 ms tones and/or noise presented at 4 bursts/s, were continuously presented in one or both ears until a single unit was encountered and isolated. Units were identified as being within the LSO based upon electrode location, recording depth, and similarity of monaural and binaural response properties to published data (Finlayson and Caspary 1991; Greene et al. 2010; Guinan et al. 1972a; Guinan et al. 1972b; Tollin and Yin 2005; Tsuchitani 1977). In particular, LSO units were identified based upon low or no spontaneous activity, and the BF of the unit with respect to surrounding areas: during insertion, electrodes tend to pass though monaurally sensitive areas before entering, and after exiting LSO (Guinan et al. 1972a; Guinan et al. 1972b; Tsuchitani 1977). LSO in the cat is S-shaped in a coronal/frontal section, thus unit-BFs increase when passing through the medial or lateral limbs, and decrease when passing thought the middle limb (Tsuchitani and Boudreau 1966). Importantly, units in LSO show ipsi-excitatory, contra-inhibitory binaural response characteristics.

When a single LSO unit was isolated, its BF and threshold were initially (qualitatively) estimated audiovisually from responses to the search stimuli. Responses to ipsilateral stimuli were then quantified using detailed frequency response maps (quantifying BF), BF-tone and noise rate-level curves (quantifying thresholds), and BF-tone post-stimulus time histograms presented at about 20 dB above threshold. BF was generally invariant with level, and was generally comparable to the characteristic frequency (CF) of the unit. Binaural sensitivity was assessed with interaural level difference (ILD) curves, created by fixing the sound level of BF-tones or noise at 10 dB above threshold in the ipsilateral ear, and increasing the sound level from 20 dB below to 20 dB above this reference level in the contralateral ear. Responses to SAM stimuli at multiple levels were recorded for ten repetitions of a BF-tone carrier 100% modulated by f_m_ logarithmically spaced from 10 Hz to over 2 kHz (or half the carrier frequency, whichever was lower).

### Data Analysis

Responses to SAM tones were quantified with average rate and synchronization modulation transfer functions (MTFs). Figure 1 shows the analysis of an example LSO unit (101013-4.02) to best-frequency (BF: 27.6 kHz; carrier) SAM tones with various modulation frequencies (f_m_). An example SAM tone with a f_m_ of 10 Hz (the lowest typically tested) and carrier frequency of 250 Hz (set well below BF for this unit for the purpose of demonstration) is shown at the bottom of the Figure 1A. The dot raster in Fig. 1A demonstrates the responses of a typical LSO unit to ten repetitions of BF-SAM tones with f_m_ increasing from 10 Hz to 2 kHz. Rate mean (black line) and standard deviation (gray area) are calculated over the entire stimulus duration (200 ms) at each f_m_, consistent with prior reports, and are shown on the same axes to the right of the raster. Synchronization (gray line) is calculated as the “vector strength” from Goldberg and Brown (1969), i.e. from period histograms constructed with 100 bins per cycle. Synchronization is only shown (in all figures) if significant via a Rayleigh test (p < 0.001). Unit sensitivity to AM stimuli is assessed as an equivalent filter shape, and are analyzed as shown for the rMTF (mean ± one standard deviation) from a second LSO unit (110321-2.02; BF 4.91 kHz) shown in figure **1B**. Rate functions were classified using tests intended to identify both *significant* and *substantial* variation across modulation frequencies. These rMTFs were classified as band-pass, low-pass, or high-pass based upon the presence of a *significant* variation across the rMTF (ANOVA, p < 0.05), and a *substantial* rate decrease to 70% of the rate observed at the best modulation frequency (BMF; a decrease of ≥ 30%, the 3 dB/half-power point) at higher and/or lower f_m_. Spontaneous activity was considered negligible and was not taken into account because unit responses tended to show very low (80% of units < 5 spikes/s) or no (34% showed 0 spikes/s) spontaneous activity, consistent with prior descriptions of LSO units in both anesthetized and decerebrate cats (Greene et al. 2010; Tsai et al. 2010). rMTFs that showed neither significant nor substantial modulation across f_m_ were classified as all-pass and are not included in most further analyses. Synchronization functions were similarly classified based on a 50% change criterion. sMTFs typically responded with band-pass or low-pass tuning. High-pass and band-reject responses were seldom encountered in either response rate or synchrony modulation transfer functions. Tuning width was quantified using the quality factor (Q; BMF/bandwidth) at 70% of the maximum value (arrows) for band-pass responses. The synchronization cutoff frequency was similarly defined as the f_m_ above the synchronization BMF at which the synchronization decreases to 70% of the maximum.

**Fig. 1.**
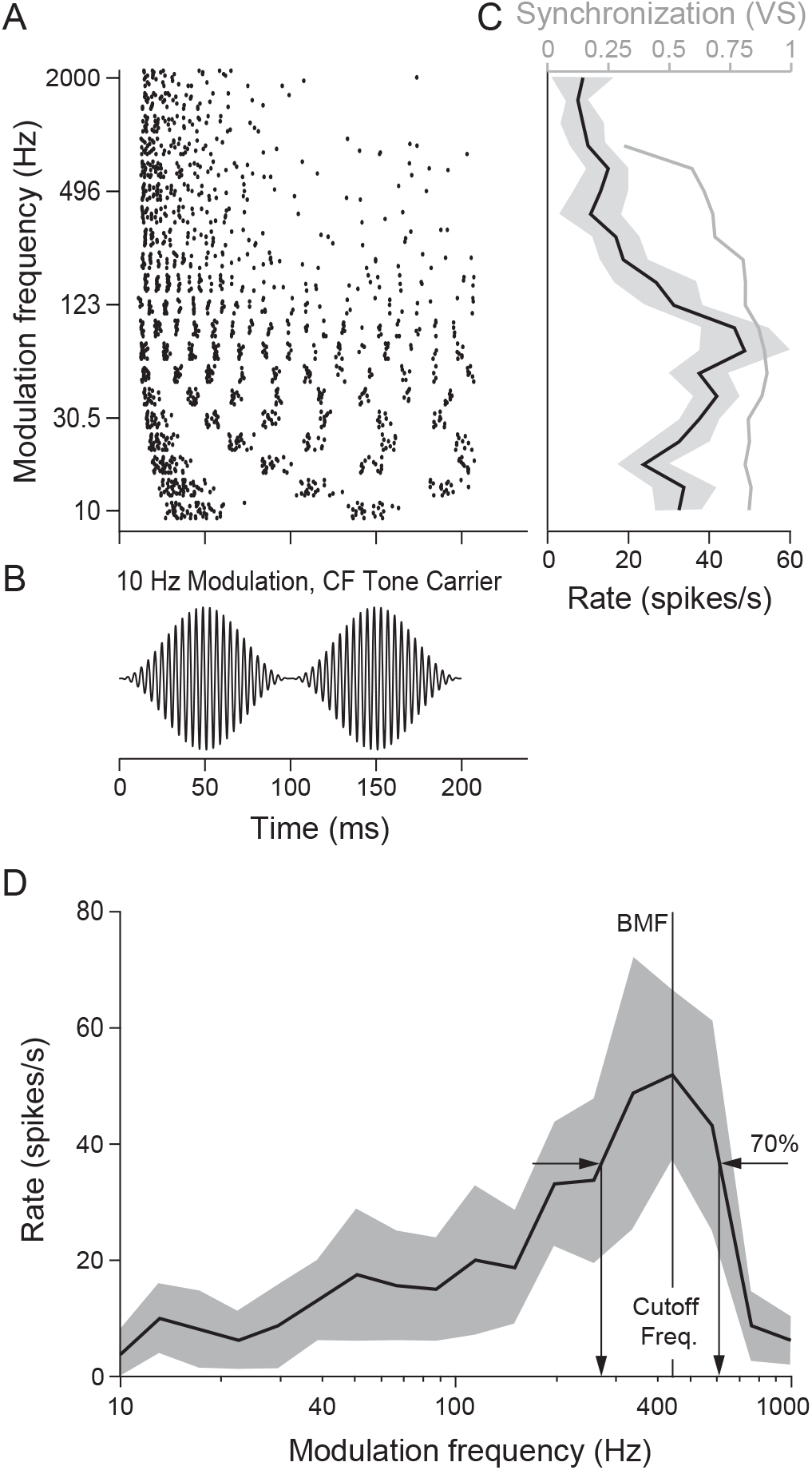
Example recordings from two single LSO units in response to SAM tone stimulation A: Responses from an example unit (101013-4.02; BF 27.6 kHz). The axes at left shows a spike raster plot for ten repetitions of SAM tones with modulation frequencies (f_m_) from 10-2000 Hz. Average rate (black) and synchronization (gray) modulation transfer functions (right; MTFs) were generated by average a unit’s responses (left) to ten repetitions of modulations from 10 Hz (bottom) to approximately 2 kHz in 20 steps. The gray shaded region indicates the mean ± standard deviation. B: rMTF from another example unit (110321-2.02; BF 4.91 kHz). Rate MTFs that show significant variation across modulation frequencies (ANOVA p<0.05) were classified based on a rate decrease to 70% of the maximum (arrows) at f_m_ above and/or below the best modulation frequency (BMF; i.e. the cutoff frequencies).

### Computational model

Details of the computational model have been published previously (Greene and Davis 2012). The responses of LSO cells were simulated in MATLAB (Mathworks, Natick, MA) using an anatomically inspired model of the brainstem ipsilateral to the LSO. An auditory nerve fiber model (Zilany and Bruce 2006; Zilany et al. 2009), provides input to the CN cells that project to LSO both directly (spherical bushy, SB) and via an inhibitory interneuron (globular bushy, GB; lateral nucleus of the trapezoid body, LNTB).

CN, LNTB, and LSO cells are modeled as single compartment integrate-and-fire neurons as described by MacGregor (1987). Each cell’s membrane capacitance (C_m_), leakage conductance (G), potassium channels (G_k_; E_k_=-10 mV) and synaptic inputs are calculated in parallel. One synaptic input is included for each population from which a cell draws input, and the sign of the associated reversal potential is set in order to determine whether the connection is excitatory (E_ex_=70 mV) or inhibitory (E_in_=-10 mV). The membrane dynamics are governed by three equations:

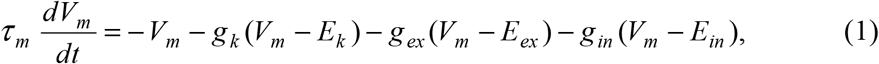

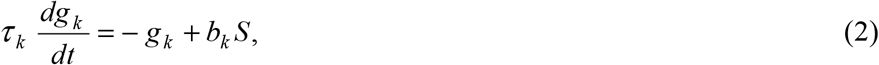

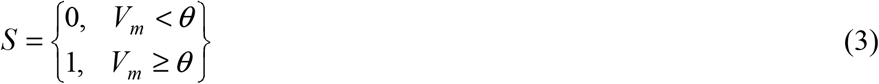

where V_m_ is the cell’s membrane potential with respect to threshold, τ_m_ = C_m_/G, g_k_ = G_k_/G, g_ex_= G_ex_/G, g_in_ = G_in_/G, b_k_ and τ_k_ are the step size and time constant, respectively, of the potassium conductance change after a spike S occurs (an absolute dead-time of 0.7ms is enforced), and θ is the cell’s threshold. Output spikes produce a step increase (δ) in the cumulative conductance of the cell’s synaptic targets (g_ex_ or g_in_) after a fixed delay, which decays exponentially with time constant, τ.

Table **1** specifies the values of both intrinsic parameters (τ_m_, θ, b_K_ and τ_K_) and connection parameters (BW, N, δ and τ), which are identical to those described previously (Greene and Davis 2012). CN and LNTB cell parameters were set to produce appropriate PST discharge patterns (primary-like, and primary-like with notch; not shown), and LSO cell parameters were set to closely match the values that produced a fast, sustained chopper response in the absence of inhibitory input (Zhou and Colburn 2010). The strength (δ_LNTB->LSO_) and delay of inhibitory LNTB input were varied to assess the effects of inhibitory input on LSO response characteristics.

**Table 1.** Model Properties. Properties describing the intrinsic (top) and network (bottom) properties of the feedforward LSO cell model. Most parameters were constrained by physiological estimates (when available), and are set to produce appropriate BF tone discharge patterns. The LSO cell potassium channel step size (bk) and time constant (τ_k_) were set to approximate values defined previously (Zhou and Colburn 2010).

| *Intrinsic* | $\tau_m$ (ms) | $\theta$ (mV) | $b_k$ | $\tau_k$ (ms) |
| --- | --- | --- | --- | --- |
| SB/GB/LNTB | 1 | 7.5 | 2 | 1 |
| LSO | 1 | 15 | 0.6369 | 5 |

Table 1. Model Properties. Properties describing the intrinsic (top) and network (bottom) properties of the feed-forward LSO cell model.
| *Connection* | $BW_{A \rightarrow B}$ (oct) | $N_{A \rightarrow B}$ | $\delta_{A \rightarrow B}$ | $\tau_{A \rightarrow B}$ (ms) |
| --- | --- | --- | --- | --- |
| ANF→SB | 0.1 | 1 | 7.5 | 1 |
| ANF→GB | 0.1 | 3 | 3 | 1 |
| SB→LSO | 0.5 | 20 | 2 | 9 |
| GB→LNTB | 0.1 | 1 | 7 | 1 |
| LNTB→LSO | 0.6 | 10 | varies | 9 |

## RESULTS

The responses to SAM tones were characterized in 44 LSO units from 13 cats. Monaural and binaural response characteristics of LSO units are comparable with previous reports in cat (Greene et al. 2010; Guinan et al. 1972a; Guinan et al. 1972b; Tollin et al. 2008; Tsuchitani 1977). BFs varied from 1 to 28 kHz similar to prior reports of cat LSO units (Greene et al. 2010; Tsuchitani and Boudreau 1966). Units are classified based on their BF-tone PST discharge pattern (i.e. Chop-S, Chop-T, Primary-like or Onset-S) when a post-stimulus time (PST) histogram was available (identified with N/A otherwise), in the manner we described previously, as we hypothesized in that prior report that this response may indicate the strength of inhibitory input to LSO cells (Greene and Davis 2012). In the following results, we first describe the responses of two units representing the range of responses observed in LSO, next we assess the basic modulation tuning characteristics of LSO cells at low sound levels (near threshold), followed by a description of the change in response characteristics at higher levels, and end with a description of the responses of a previously developed computational model to SAM tone stimuli.

Figure 2 shows the responses of two typical LSO units to BF pure tones as a function of level, and BF SAM tones at several levels as a function of the modulation frequency (A-C: unit 101013-4.10, BF 17.6 kHz; D-F: unit 101013-4.01, BF 20.3 kHz). Figs. 2A and D show BF-tone rate-level curves, and demonstrate the monotonically increasing response rates with increasing sound level above threshold typical of LSO unit responses in decerebrate cat (Greene et al. 2010). The responses of these units to AM stimuli were assessed at multiple levels, which are indicated in 2A and 2D as gray-shaded dots (black-low, gray-high). Gray shaded lines in Fig. 2B/E and 2C/F represent rMTFs and sMTFs, respectively, recorded at the levels indicated by the corresponding gray shaded circle in 2A and 2D. Similarly, the response rate elicited by a pure tone at the same sound level, as indicated by the gray shaded circle, is shown on the y-axis in 2B and 2E. Near threshold (black, ~10 dB above or below), the rMTFs show band-pass responses: discharge rates either matched (2B) or showed somewhat lower (2E) response rates at low and high f_m_ than were elicited by pure tones, and increase to distinct peaks that tended to show rates somewhat higher than the pure tone rates. At higher sound levels the response rates at low and high f_m_ increase disproportionately more than the rate at the BMF, thus the rate variation across f_m_ decreases and the rMTF could show all-pass tuning (such as in 2E), retain more weakly varied band-pass, or show a more complex response such as the band-reject response in 2B. The rMTFs of the two representative units shown thus reveal a trend of well-tuned, band-pass responses at low, and poorly-tuned responses that approach all-pass responses at higher sound levels.

**Fig. 2.**
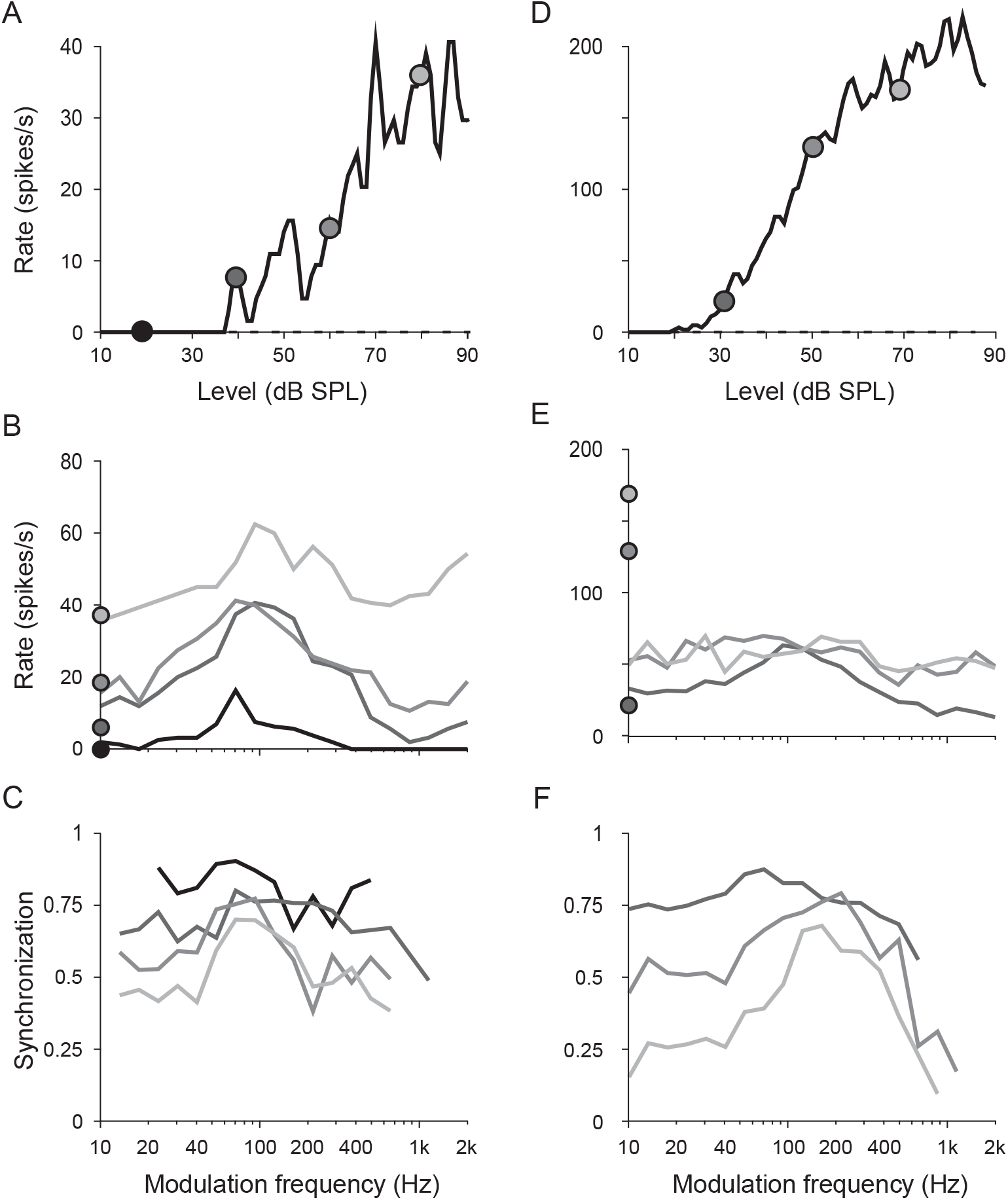
Responses to BF and SAM tones as a function of level. The responses of two typical LSO units (A-C: 101013-4.10 BF 17.6 kHz; D-F: 101013-4.01 BF 20.3 kHz) to BF-tones (A/D) and SAM-tones (B-C/E-F) reveal discharge rates that increase and synchronization that decreases with increasing sound level. rMTF shape typically shifts from band-to all-pass, and sMTF shape typically shifts from low-to band-pass with increasing level. Circles in A/D and gray shade in B-C/E-F indicate the level at which the SAM stimulus was presented, and circles in B/E show the corresponding BF-tone rate.

LSO unit synchronization to the SAM tone envelope, in contrast, typically shows an opposite trend with sound level as the rate functions. As demonstrated in 2C and 2E, sMTFs are generally maximal at low sound levels, and typically show low-pass responses (note that decreasing synchronization is not always visible because we only show synchronization when significant). Synchronization generally decreases at all frequencies with increasing sound level; however, sMTFs often decrease substantially at low, decrease only slightly at intermediate, and remain low at high f_m_ (above the synchronization cutoff frequency, defined as the 3dB point), thus sMTFs appear more strongly modulated and show band-pass tuning at high sound levels. Note that synchronization is not shown for f_m_ at which the response did not significantly synchronize to the stimulus, where synchronization is typically low. The sMTFs shown once again represent the range of responses observed in, demonstrating a clear trend from low-to band-pass tuning in many units (Fig. 2F), and a more variable, less distinct trend in a minority of units (Fig. 2C).

### MTFs at low sound levels

As demonstrated by the examples shown in Fig. 2, LSO units typically show the sharpest rMTF tuning, and the highest overall synchronization at sound levels near (within ~30 dB) their BF tone threshold. In the following analyses, therefore, rMTFs and sMTFs are characterized at the recorded level eliciting maximum synchronization at any frequency.

The discharge rates recorded at the lowest f_m_ were correlated with the rates to pure tones at the same level (R^2^= 0.627, p ≪ 0.001), and many units show very similar response rates, as demonstrated in Fig. 2B; however across the population of units sampled, median rates were significantly lower for SAM tones at the lowest f_m_ tested than to pure tone (24.4 vs 40 spikes/s, Mann-Whitney U test, p = 0.03). An example of a large disparity in discharge rate is demonstrated by the unit in Fig. 2E, showing a ~3/1 difference between the pure tone and SAM tone responses. In contrast, maximum response elicited by SAM tone stimuli (i.e. the mean rate elicited at the rBMF) tended to show somewhat higher rates than a pure tone at the same sound level, though medians were not significantly different (40 vs 49 spikes/s, Mann-Whitney U test, p = 0.5), though they remained strongly correlated with the tone discharge rate. Discharge rates, therefore, appear to be reasonably well characterized by a modulation of the rate elicited by a pure tone.

The best modulation frequency varies across a broad range of f_m_ for LSO units that have band-pass rMTFs. Fig. 3A shows rMTFs from four representative units with undefined (dark gray; unit: 101013-4.01, BF: 20.3, BMF: 0.093), Chop-S (black; unit: 110607-3.02, BF: 9.29, BMF: 0.26; and unit: 101216-1.04, BF: 10.3, BMF: 0.70) and Primary-like (light gray; unit: 110413-1.05, BF: 8.8, BMF: 2.19) PST discharge patterns, and are shown normalized to their maximum (i.e. rate at BMF); spontaneous rates were essentially zero (i.e. < 0.25 spikes/s) in all four units. In each case, the discharge rate at each f_m_ showed some variability, but for clarity error bars are not shown. Fig. 3B shows the distributions of rate BMFs for band-pass responses of each discharge pattern type, which were identified using our previously described PST classification scheme (Greene and Davis 2012). Units for which a PST was not collected, and a discharge pattern not defined, are represented by white bars and labeled as not available (N/A). rBMFs determined in 30 of 32 (94%) units were between 20 and 2000 Hz, with a median of 273 Hz. Distributions of units grouped by PST discharge types overlap considerably, and are not significantly different when assessed with a one-way analysis of variance (ANOVA; F_3,39_= 0.82, p = 0.489). Likewise, rBMF appears generally independent of a unit’s BF, showing little or no correlation between the two measures (R^2^= 0.001, p = 0.828).

**Fig. 3.**
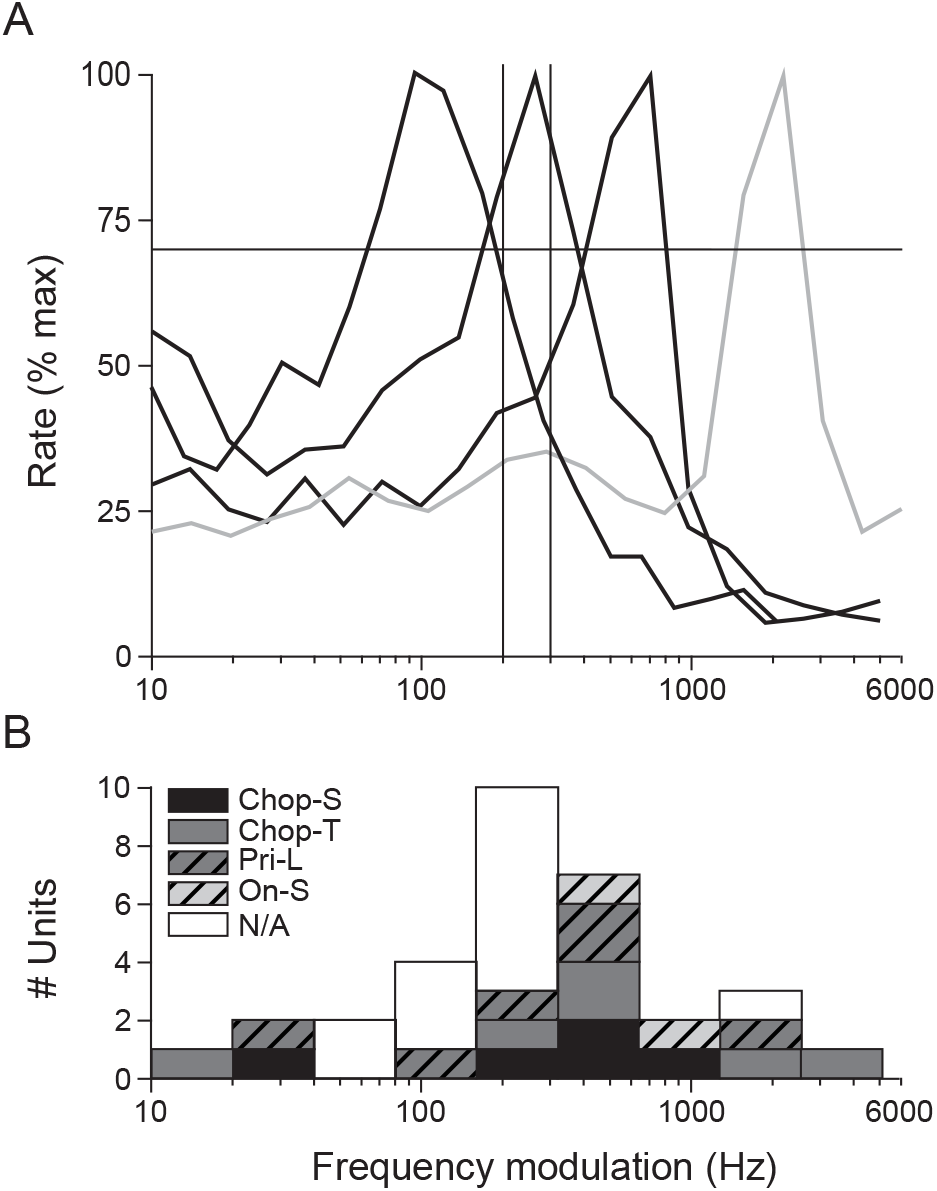
BMF range. A: rMTFs from four typical LSO units demonstrate the range of BMFs recorded (unit, BF (kHz), BMF (kHz): 101013-4.06, 19.2, 0.04; 110607-3.02, 9.29, 0.26; 101216-1.04, 10.3 0.70; 110413-1.05, 8.8, 2.19). B: Histogram of BMFs from all band-pass units. Responses are segregated based on discharge patterns observed in BF tone PST histograms. White bars indicate units for which a PST was not available (N/A), thus discharge pattern was not defined.

Figure 4 shows the sharpness of tuning of rMTF curves as a function of rBMF for units that have band-pass rMTFs. Sharpness of tuning is quantified for each unit with the quality factor (Q), defined as BMF / bandwidth at 70% of the max (e.g. Fig. 1B). Unit tuning show values below 2, comparable to reports in ICC from anesthetized cat (Langner and Schreiner 1988) and awake rabbit (Nelson and Carney 2007), though with substantially higher rBMFs. Tuning sharpness across the population of units sampled appears independent of BMF (correlation R^2^= 0.128, p = 0.715), and no significant variation across discharge pattern types is evident when assessed with a one-way ANOVA (F_4,38_= 0.65, p = 0.63). Taken together, these data suggest that rate tuning characteristics of LSO units are generally independent of both their pure tone frequency sensitivity and their PST discharge characteristics.

**Fig. 4.**
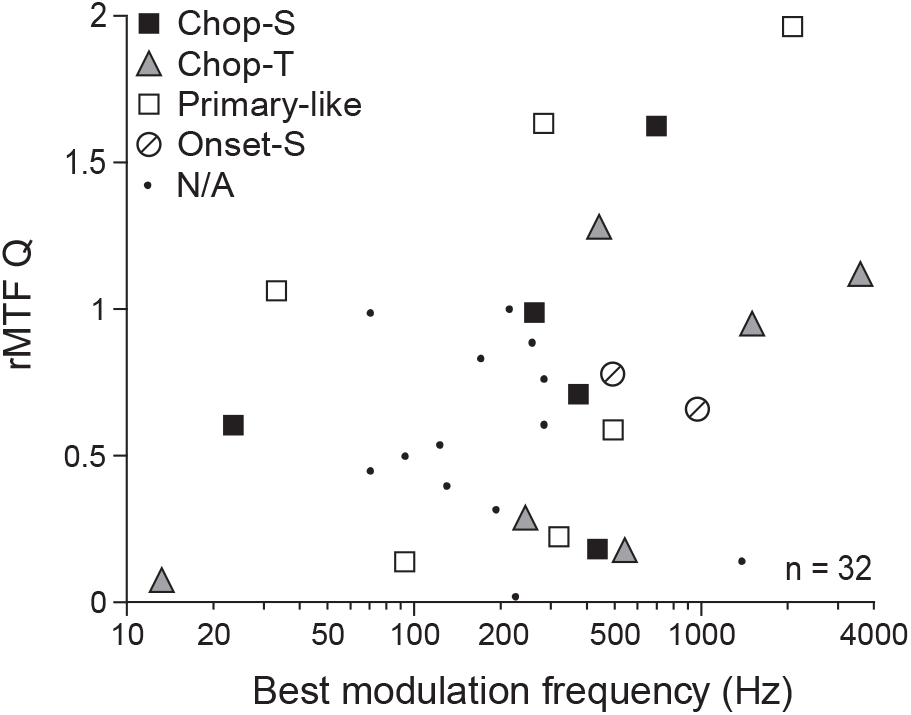
SAM-tone tuning width. rMTF tuning width, shown as the Q value (BMF/bandwidth), is shown as a function of BMF for band-pass responses, and does not vary across discharge patterns. Gray shaded area indicates the range of responses observed in awake rabbit ICC (Nelson and Carney 2007). Symbols indicate BF-tone PST discharge patterns.

Figure 5 shows the maximum synchronization elicited (value at sBMF) as a function of the synchronization cutoff frequency. A single unit showed an all-pass response, thus was excluded from further analysis. LSO units synchronize to the modulation relatively well: all but one unit show vector strengths greater than 0.7, and most (34/45) respond with greater synchronization than the best values observed in the auditory nerve and spherical bushy cells (horizontal line; Joris and Yin 1998; Joris and Yin 1992). Primary-like units tended to show the lowest strengths at each synchronization cut-off frequency, and Onset-S units tended to show the highest synchronization cut-off frequencies; however, synchronization strength is not significantly different across discharge pattern types when assessed with a one-way ANOVA, (F3,24 = 1.36, p = 0.28).

**Fig. 5.**
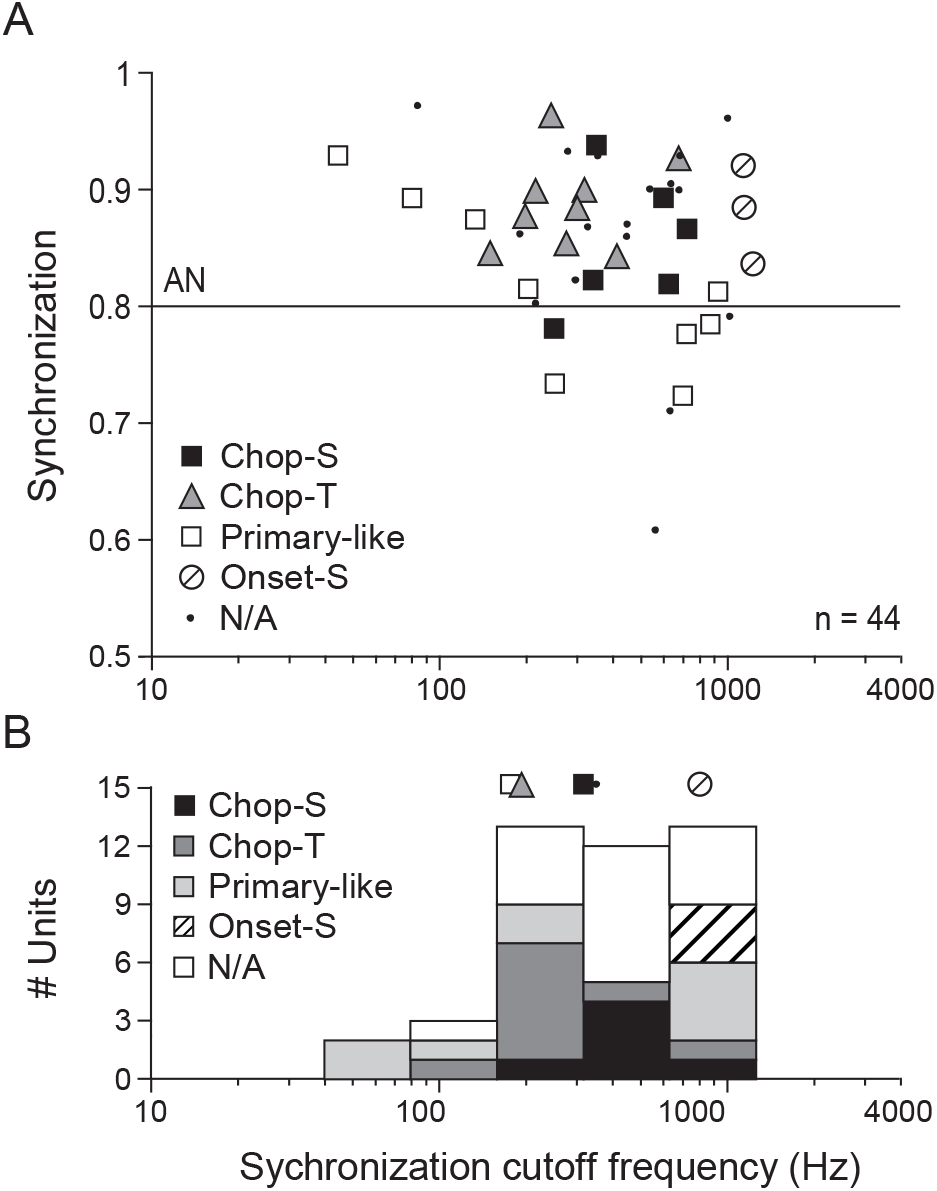
Maximum synchronization. A: The maximum synchronization elicited (at any level) is shown as a function of the synchronization cutoff frequency (at the same level) for all units, and is generally higher than values observed in the auditory nerve. B: Histogram of synchronization cutoff frequencies. Symbols and bar shades indicate BF-tone PST discharge patterns. Maximum synchronization of the auditory nerve is from Joris and Yin (1998; 1992).

A histogram showing the synchronization cutoff frequency (i.e. the f_m_ at which the synchronization has dropped by 3 dB) is shown for each PST response type in Figure 5B. Symbols along the top of the plot indicate median values for each discharge pattern type. A one-way ANOVA (log2 transformed) indicates a significant difference in the distributions (F_3,24_= 3.40, p = 0.034), and Bonferroni corrected pair-wise comparisons indicate that the Onset-sustained units show higher cutoffs than other unit types (students t-test p < 0.05). In contrast to rate, therefore, synchronization to the AM envelope appears somewhat related to PST discharge type, thus may indicate some characteristics of an inhibitory input to the LSO cell.

### MTF tuning as a function of level

The trend with level observed in the examples shown in figure 2 are generally typical of the sample of units recorded. That is, discharge rate tends to increase, rMTF tuning decreases, synchronization strength decreases, and sMTF sharpness of tuning increases at higher sound levels. The rMTF response types for all LSO units recorded are shown in figure 6A as a function of sound level above threshold. Similarly, the proportion of units showing each response type are shown as a function of level in figure 6B. Below the BF-tone threshold (0 dB), responses to AM stimuli are scarce and appear equally likely to produce band-pass and all-pass responses. Within the dynamic range of the BF-tone rate-level curve (0 - ~30 dB), band-pass responses predominate, and the proportion of all-pass responses slowly increases. At higher levels (near the level at which the response to a BF-tone typically saturates, all-pass responses predominate, although a minority of LSO units could respond with rMTFs with any response type at most sound levels. Anecdotally, onset-sustained responses, tend to maintain their band-pass responses except at the highest sound levels recorded (50-70 dB above threshold; not shown).

**Fig. 6.**
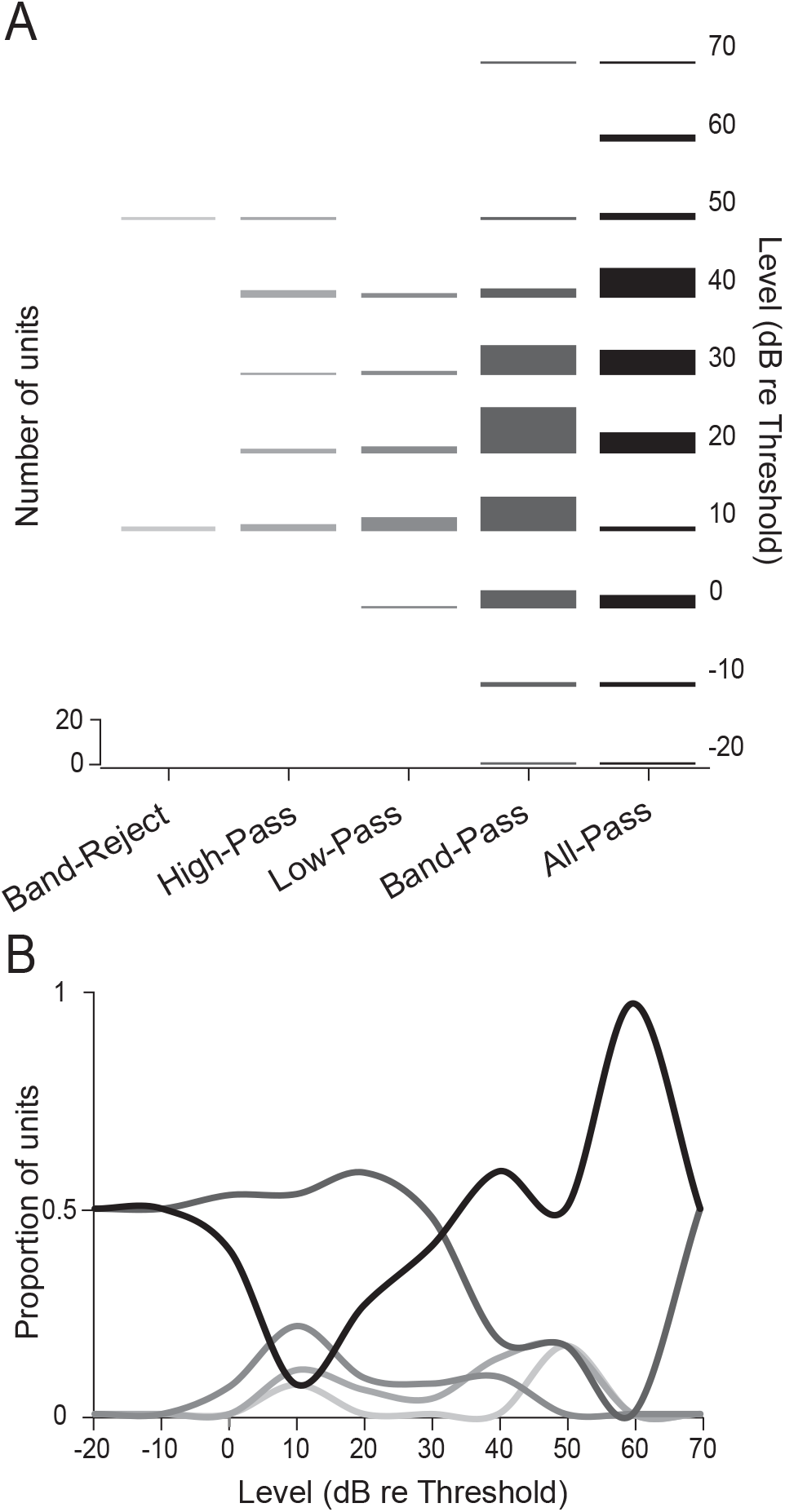
rMTF shape as a function of stimulus level. A: Histograms of rate tuning shape are shown in 10 dB bins with respect to the stimulus level above the unit’s BF-tone threshold, and reveal a shift from band-to all-pass response shapes. B: Proportion of units showing each tuning type as a function of level above the BF-tone threshold. Gray shades correspond to those in the bar graph shown in A.

sMTF filter types are shown as a function of sound level above threshold in figure 7A for all LSO units recorded, and the proportion of units responding with each synchronization response type are shown as a function of sound level above the BF-tone threshold in figure 7B. Synchronization tends to drop off at high f_m_ at all sound levels, thus low- and band-pass responses predominate at all sound levels. As demonstrated in the Fig. 2, synchronization tended to decrease more strongly at higher and lower than intermediate f_m_ with increasing sound level in most units. Consistent with these examples, response types shift from low-pass near threshold, to band-pass responses above ~30dB re threshold, near the level at which responses to BF-tones saturate. In contrast to rMTFs, synchronization functions are generally similar, showing low-pass sMTFs at low sound levels, band-pass functions at higher levels with few differing responses.

**Fig. 7.**
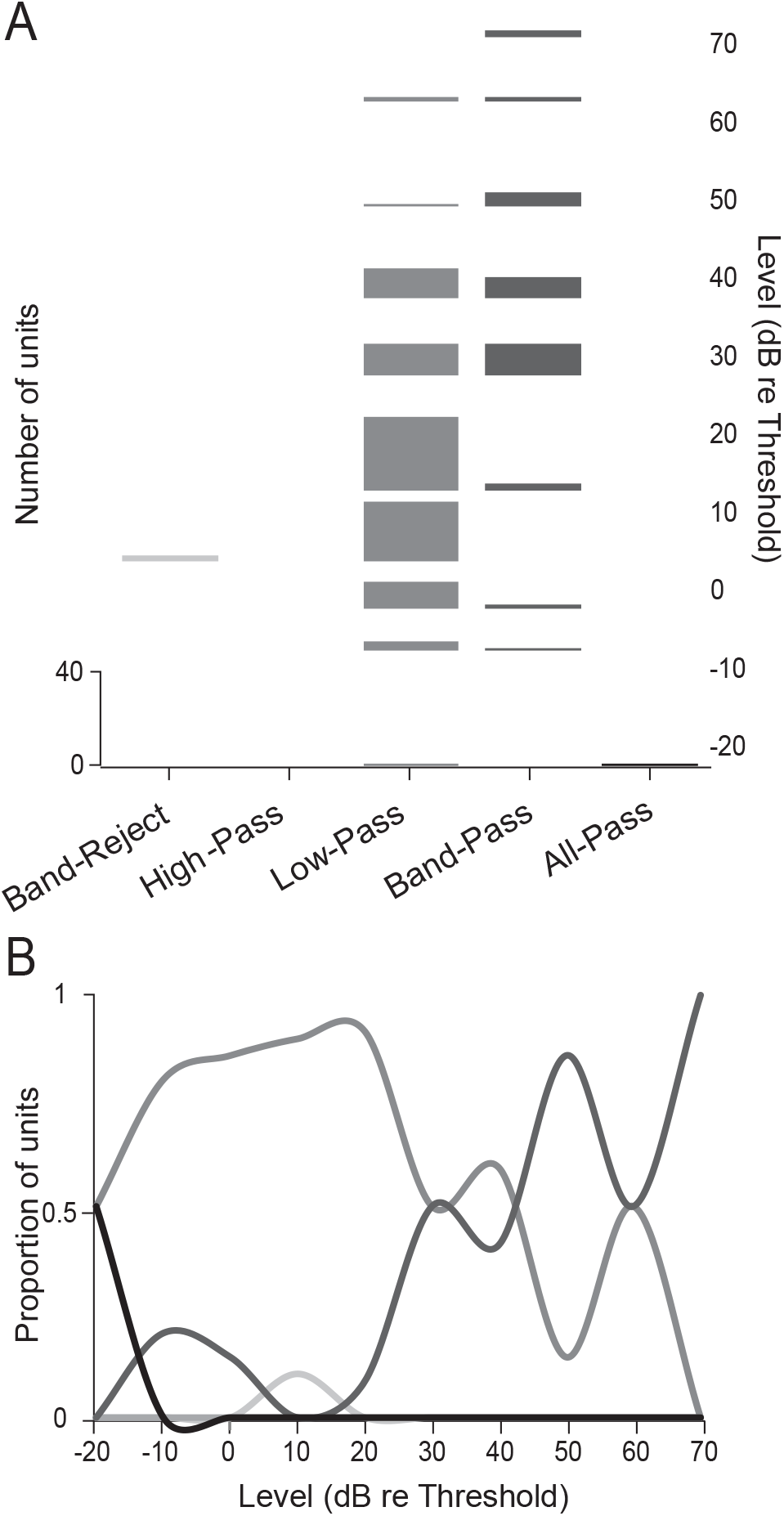
sMTF shape as a function of stimulus level. A: Histograms of synchronization tuning shape are shown in 10 dB bins with respect to the stimulus level above the unit’s BF-tone threshold. B: Proportion of units showing each tuning type as a function of level above the BF-tone threshold. Gray shades correspond to those in the bar graph shown in A.

### Model LSO responses to SAM stimuli

Our hypothesis is that the more clearly defined band-pass tuning in decerebrate cat, compared to the mostly low-pass responses recorded in barbiturate anesthetized cat, mirrors the lower regularity observed in response to pure tone PST histograms in decerebrate compared to anesthetized cat, thus may result from the action of an anesthetic dependent blockade of glycinergic inhibition. That is, anesthesia could act to reduce the effectiveness of inhibition in LSO, thus producing poorer rate tuning. In order to evaluate this possibility, we presented SAM stimuli to a computational model of LSO developed to explain the differences observed in PST histograms to BF tone stimulation (Greene and Davis 2012). The model organization is shown in figure 8A, and is identical to our prior report (Greene and Davis 2012) except for an update to the auditory nerve model (Zilany et al. 2009). Responses of the model LSO cell to SAM tones with varying inhibitory input strength is shown in Fig. 8B. Comparable to the excitatory bushy cell input, the response to a 40 dB SPL SAM tone shows an all-pass rMTF when inhibition is weak. Increasing strength suppresses responses at all f_m_, but particularly so at low and high f_m_ resulting in band-pass rMTF shapes when inhibition is strong, while intermediate inhibitory strength can produce a low-pass shape. The inhibitory input is not delayed with respect to the excitatory input in order to generate Chop-S, Pri-like, and Onset_S type responses with increasing inhibitory strength (Greene and Davis 2012).

**Fig. 8.**
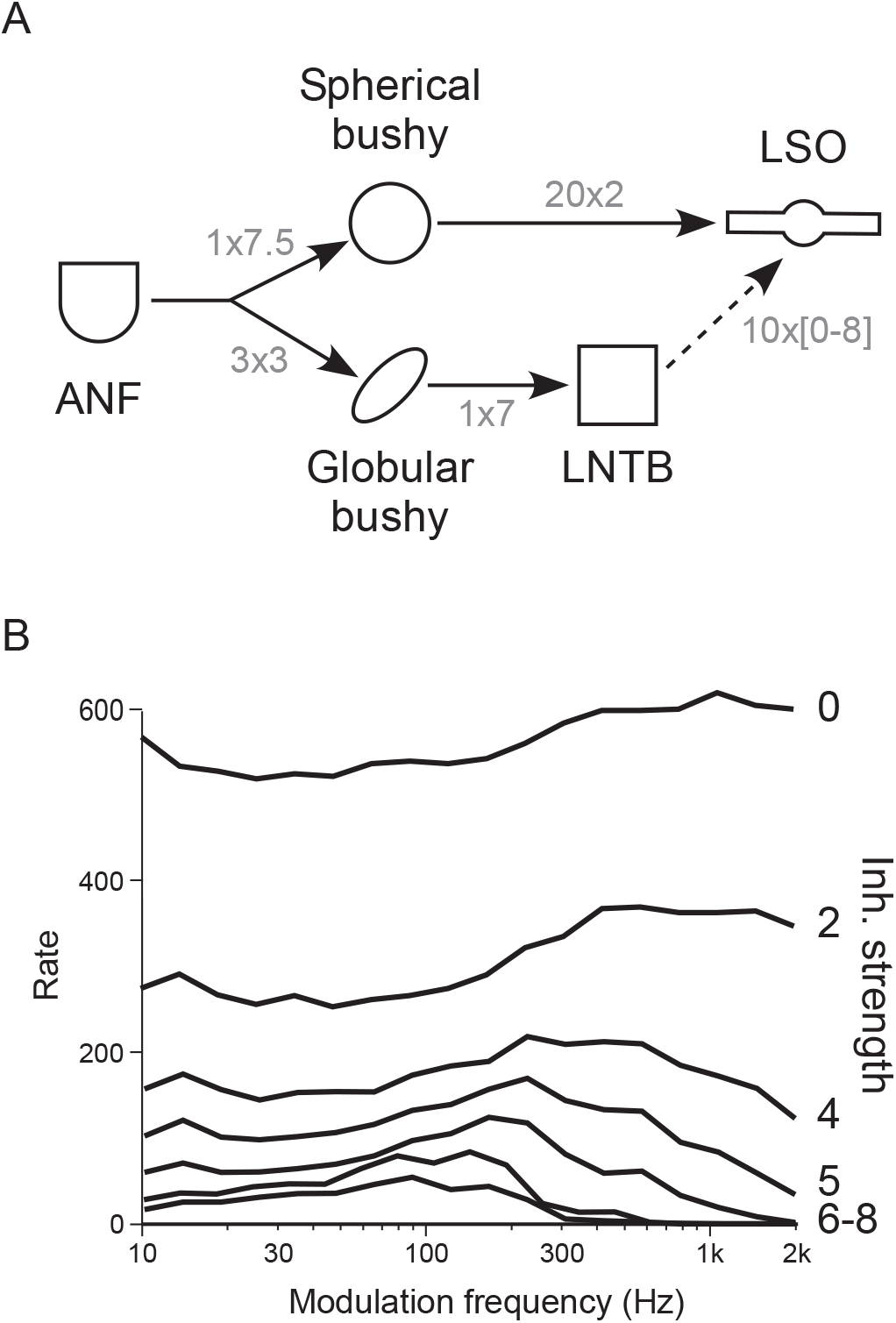
LSO model organization. A: Organization of the feed-forward inhibition LSO model. Lines indicate connections between unit groups (symbols). Numbers indicate number of projections converging on each unit x strength of that connection. B: Response of the LSO model with strength of LNTB (Inh.) input from 0 to 8. Inhibitory delay = 4ms. AM stimulus presented at 40 dB SPL.

In order to assess the effects of inhibitory strength on the responses of the model to SAM stimuli with varying sound levels, responses of the model elicited by three sound levels are shown in figure 9 for a range of inhibitory strengths. Model rMTFs and sMTFs are displayed in the left and right columns, respectively, such that inhibitory strength increases in rows from top (0) to bottom (6). Three sound levels, from low (near the model unit’s BF tone threshold; black) to high (a level above which the model response has saturated to a BF-tone rate-level function; light gray) are shown in each plot (i.e. stimuli were presented to the auditory nerve model at 20, 40, and 60 dB SPL). When the inhibitory strength is zero the average rate increases monotonically with sound level and shows little variation across f_m_, while synchronization shows low-pass responses at low, and rapidly decreasing synchronization at higher sound levels. The effects of low but non-zero inhibitory strength (2) manifest as band-pass sMTFs at the intermediate sound level and little change in rMTF modulation despite substantially decreased discharge rates. At a more intense inhibitory strengths (4), which would produce Pri-like or Onset-S type responses in BF tone PST histograms (Greene and Davis 2012), rMTFs show a more prominent peak at the low sound level, with a low-pass sMTF, and all-pass or a more complex rMTF and sharply tuned band-pass sMTFs at higher sound levels. At a high inhibitory strengths (6), where response rates to pure tone PST histograms are substantially suppressed, rMTFs are band-pass and sMTFs are complex or all-pass at the intermediate and high sound levels.

**Fig. 9.**
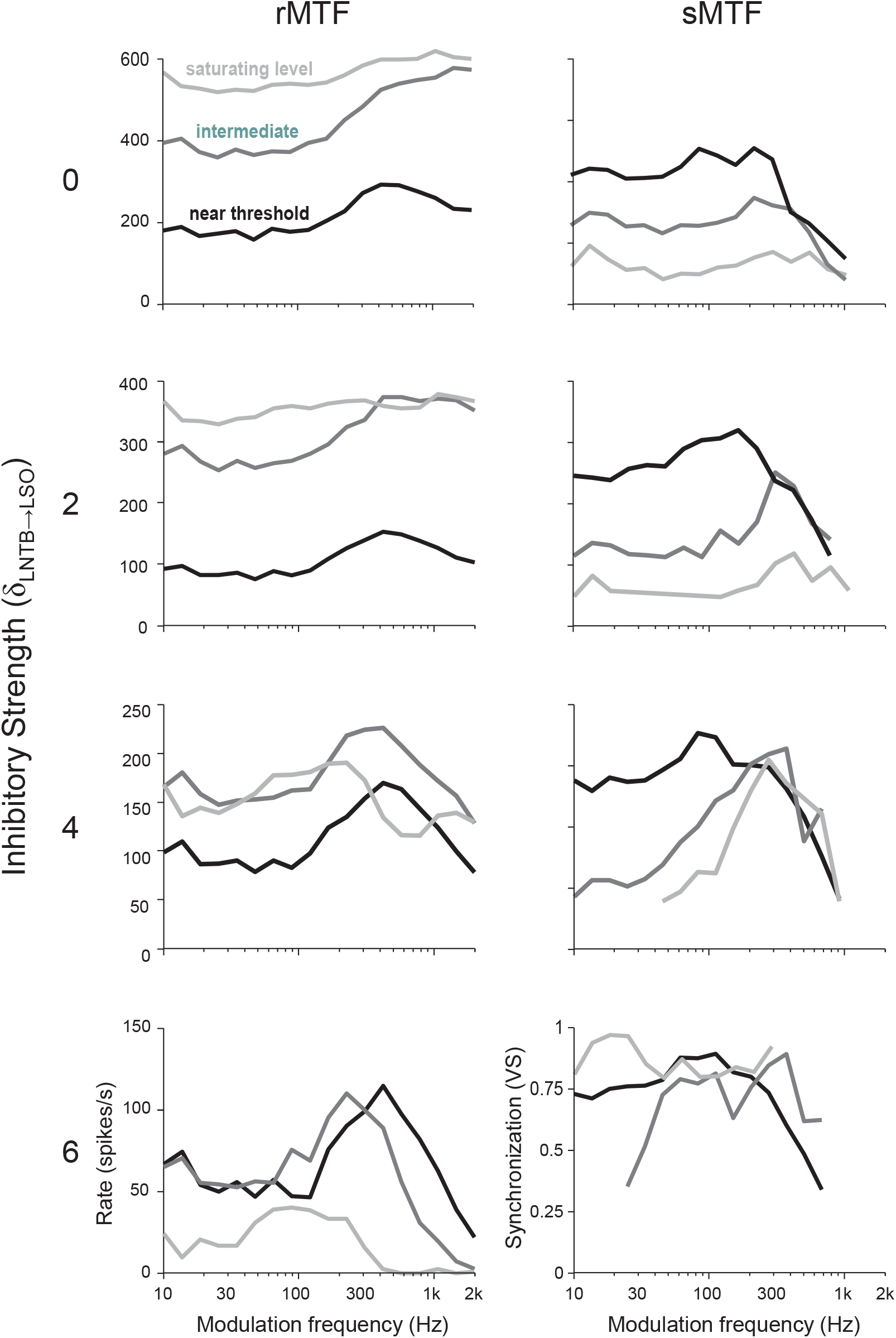
Model LSO unit responses with stimulus level. Moderate inhibition reproduces features of in vivo LSO responses. Model LSO rMTFs (left) and sMTFs (right) with LNTB strength 0 (A), 2 (B) 4 (C), and 6 (D). Responses are shown for SAM tones presented at 20 (black), 40 (dark gray) and 60 (light gray) dB SPL.

Response rate increases monotonically in response to increasing sound levels for weak to moderate inhibitory strengths. When inhibition is strong, however, response rates tend to first increase then decrease dramatically with increasing sound level, although the response never becomes completely inhibited. Conversely, maximum synchronization, which is generally maximal at the lowest sound level presented, is typically relatively poor (< 0.75) when inhibition is weak, increases with increasing inhibitory strength, and nearly reaches 1 when inhibition is strong. Rate and synchronization, therefore, appear inversely correlated, consistent with the example units observed in Fig. 2. Responses of the model thus reproduce the general trends with level observed in the LSO, when the inhibitory input is sufficiently strong (≥ 4), comparable to the range of inhibitory strengths required to elicit appropriate PST discharge patterns described previously (Greene and Davis 2012).

Previously, we showed that the timing of the inhibitory input with respect to the excitatory input (in addition to the strength) affected the resulting PST histogram discharge pattern (Greene and Davis 2012). In order to assess the effect of input timing on the responses to SAM tone presentation, responses of the model with inhibitory delays of −2 ms (inhibitory inputs precede excitatory), +2 ms, and +4 ms (inhibitory input is delayed compared to excitatory) are shown in Figure 10 compared to responses of the cell with no delay. Responses were assessed for a high inhibitory strength (6) and a saturating sound level (60 dB SPL). Response rates were somewhat dependent upon inhibitory delay, with the maximum observed discharge rate (i.e. the rate at rBMF) observed varying between ~30 and ~70 spikes/s (discharge rate was lowest with delays of 0 or +2 ms). Interestingly, while responses to high f_m_ SAM tones were strongly suppressed for all inhibitory delays, responses rates to low f_m_ were more variable, showing a 3/1 ratio between the rate elicited at the rBMF and the lowest f_m_ assessed for −2 and +4 ms delays, and approaching a 1/1 ratio for a +2 ms delay. Synchronization strength likewise showed very similar trends across inhibitory delays, and in all cases tended to show low-pass responses for low- and band-pass responses for high-intensity stimuli. In all four conditions, model rMTFs reproduced the band-pass to all-pass transition with increasing sound level (not shown). An interesting effect of the fixed inhibitory delay is observable as reduced synchronization in the −2 ms and +4 ms delay conditions at frequencies near 1/delay (i.e. near 250ms and 500ms here). Neither inhibitory strength nor timing substantially affected the BMF in both rate and synchronization functions.

**Fig. 10.**
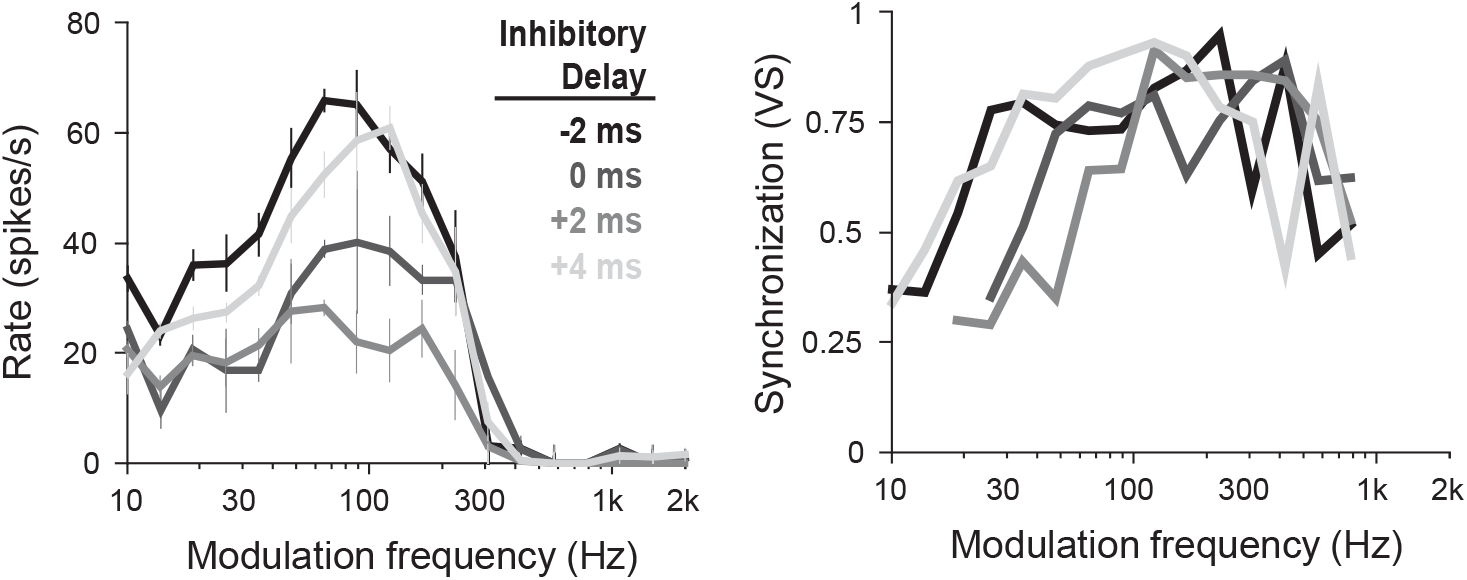
Model LSO unit responses as a function of inhibitory delay. rMTFs (A) and sMTFs (B) for an LSO cell with strong inhibitory inputs (6), stimulated with high level (60 dB) SAM tones. Inhibitory delay modulates affects the discharge rates elicited, and the rate dependence on f_m_.

## DISCUSSION

In this study we set out to test the hypothesis that single units in the LSO of unanesthetized, decerebrate cat show enhanced rate tuning to SAM tones compared to cells recorded in barbiturate anesthetized cat, consistent with the effects of an anesthetic dependent blockade of an ipsilaterally driven inhibitory input. Above, we presented evidence suggesting that LSO in decerebrate cat do indeed show band-pass responses in rMTFs at low sound levels, though that tuning is lost at higher sound levels. We went on to show that a computational model of LSO with an ipsilaterally driven inhibitory input can indeed reproduce the range of responses observed in-vivo, suggesting this is a plausible mechanism underlying these results. In the following discussion, we expound upon the differences noted between decerebrate and barbiturate anesthetized cat, the implications for these results on the likely mechanisms producing tuned rMTF functions, and speculate on the role of the LSO in processing amplitude modulated stimuli.

### Rate tuning in the decerebrate cat LSO

The responses of LSO units to SAM tones in the cat were previously described in thirteen units from a single study utilizing barbiturate anesthesia. The rMTFs of these units were collected by digitally tracing the responses of units presented by Joris and Yin (1998), and are plotted on a semi-log scale in figure 11A to allow direct comparisons to the current data. Responses are shown with a gray shade corresponding to the calculated rMTF filter shape (i.e. black all-pass, dark gray: low-pass, and light gray: band-pass). In contrast to the well-tuned rMTFs recorded in many decerebrate cat LSO units (e.g. Fig. 2C), these units show either weakly tuned low-pass or untuned all-pass rMTFs. In order to compare as directly as possible to these data we assessed responses at the level eliciting maximum synchronization and a minimum discharge rate (27 spikes/s; the lowest peak rate in the anesthetized data), which was approximately 10 dB above threshold for most units. Tuning of rMTFs was characterized in the same manner for the anesthetized and the current data, i.e. using a reduction to 70% of the max and assessed from 40 Hz to 2 kHz f_m_, and are compared in the bar plots shown in figure 11B.

**Fig. 11.**
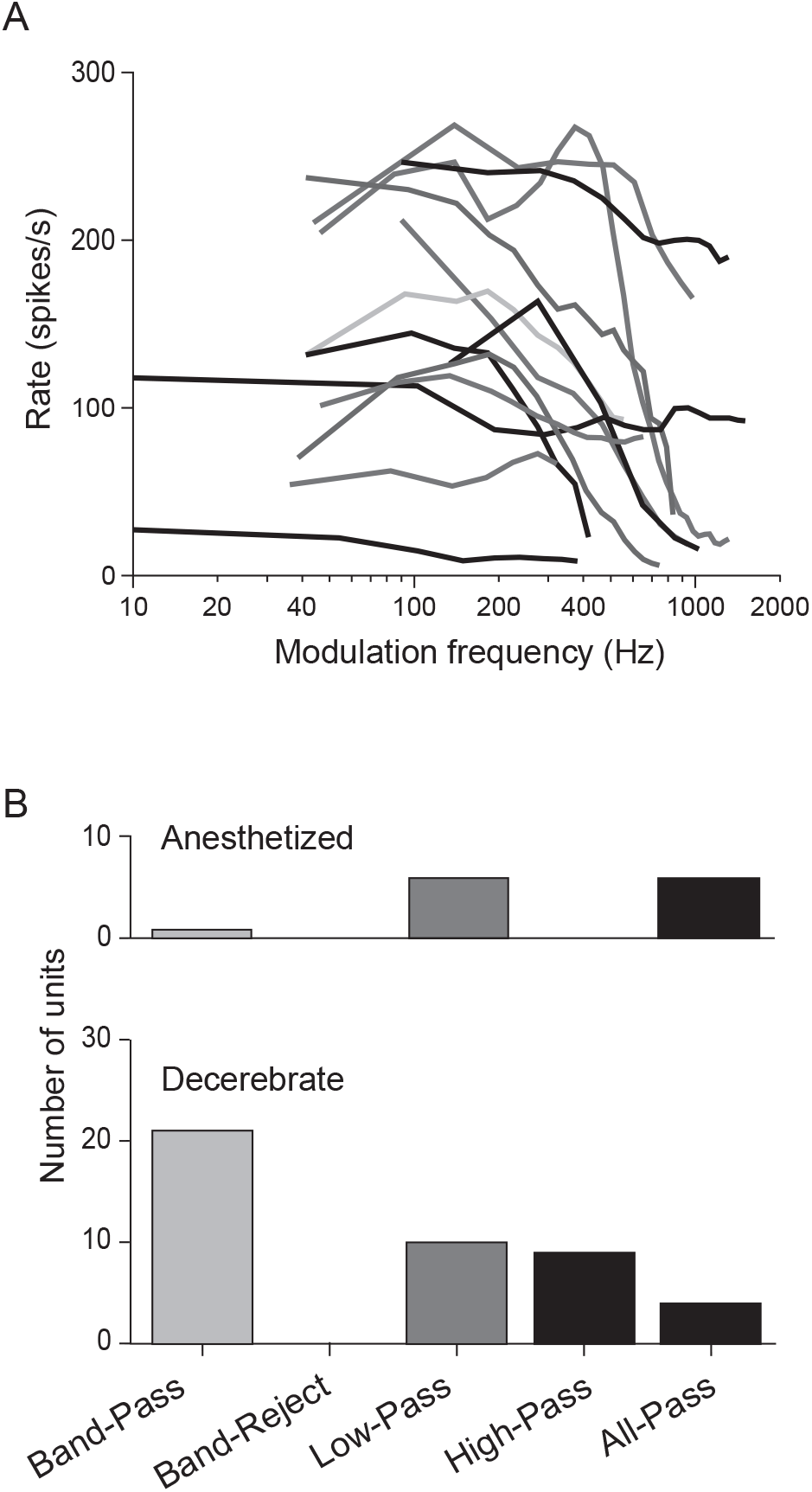
Comparison of rMTF tuning in the LSO of anesthetized and decerebrate cats. rMTFs from anesthetized (A-B) and decerebrate (C-E) cat reveal different tuning characteristics (F). Gray-shading indicates the rMTF shape determination (black: all-pass, dark gray: low-pass, light gray: band-pass). Anesthetized cat data is derived from (Fig. 13B from: Joris and Yin 1998).

Units recorded in anesthetized cat show equal numbers of low- or all-pass responses, while band-pass responses predominate in decerebrate units, with smaller proportions of low-, high-, and all-pass responses; only one unit (light gray) recorded in anesthetized cat curve is classified as band-pass in the current classification scheme. One difference that could explain this disparity between the two samples could be in the f_m_ surveyed. While the current data were typically sampled down to 10 Hz, units recorded in the anesthetized preparation were collected down to only 50 Hz modulation. Anesthetized responses, therefore, may have been insufficiently sampled to identify the low-f_m_ rate decrease required for band-pass classification, and were, therefore, instead labeled as low-pass. However, while insufficient f_m_ range could explain the larger number of low-pass rMTFs, it cannot explain the increased prevalence of all-pass units in anesthetized compared to decerebrate cat LSO (~ 50% all-pass in anesthetized vs < 10% in decerebrate cat).

The preponderance of well-tuned rate functions (at least for low sound levels) in the decerebrate cat, therefore, represents an anesthetic dependent difference in response characteristic between the two preparations. Nevertheless, the proportion of low-pass responses in the anesthetized cat LSO still represents a larger change from their inputs, the spherical bushy cells of the anteroventral cochlear nucleus, than we would have expected from reducing inhibitory strength in our LSO model. This difference suggests that either inhibition is incompletely silenced by surgical levels of barbiturate anesthesia, or that an additional mechanism is affecting unit responses.

### On the mechanism producing a modulated discharge rate

In an attempt to explain the low-pass responses recorded in anesthetized cat LSO, Joris and Yin proposed that a strong spontaneous inhibitory input from medial nucleus of the trapezoid body (MNTB), which is highly spontaneously active even in barbiturate anesthetized preparations (e.g. Koka and Tollin 2014; 0-150 spikes/sec., mean of ~26 spikes/sec.), could produce this effect (Joris and Yin 1998). However, a recent modeling study (Wang and Colburn 2012) concluded, after investigating this hypothesis, that an unrealistically high number of inhibitory inputs (each firing at 30 spikes/sec.) is required to reproduce the rMTF tuning observed. Wang and Colburn instead proposed that a low-threshold potassium channel (K_LT_) was the more likely mechanism modulating rate functions. However, we believe that K_LT_ is unlikely to be the only, or dominant, mechanism producing band-pass rate responses to SAM tones in most LSO units for two reasons. First, K_LT_ is expressed in a gradient along the tonotopic axis, where high frequency units are the least likely to express, and show the lowest expression density of K_LT_ (Barnes-Davies et al. 2004). Second, the effects observed require an anesthetic dependent mechanism. Direct evidence for such an effect has been observed previously in LSO (Wu and Kelly 1994), and for glycinergic inhibition more generally (Lu and Xu 2002), but not K_LT_ (to the best knowledge of the authors). Interestingly, modulation of an inhibitory input could affect K_LT_ indirectly, since modulation of inhibitory inputs can enhance the effectiveness of K_LT_ to periodic stimuli by promoting post-inhibitory facilitation of spiking via transient inactivation of the K_LT_ channel (Dodla et al. 2006), but this mechanism appears inadequate to explain the effects observed in decerebrate vs barbiturate anesthetized cat.

In order to address these shortcomings, we propose a similar model to Wang and Colburn (2012) but with an alternate source of inhibitory drive to LSO cells. We have previously presented physiological evidence suggesting that an additional, previously un-accounted for, ipsilaterally driven inhibitory input modulates responses in the unanesthetized decerebrate cat LSO (Greene and Davis 2012). Ample evidence suggests that spontaneously active inhibitory inputs arrive in LSO from MNTB (Koka and Tollin 2014); however, the modeling study by Wang and Colburn (2012) appeared to rule out a spontaneous source of inhibitory input, since an unrealistically large number of inputs was required in order to generate tuned rMTFs in the LSO. The underlying assumptions and organization of our model differ from those presented by Wang and Colburn (2012) in two important ways. First, rMTFs are more often and more strongly band-pass tuned than in anesthetized cat, when compared at similar sound levels. Second, since we propose an ipsilaterally driven source of inhibition to LSO, which can provide a dramatically larger number of inhibitory potentials than a spontaneous source, a few high-rate driven inputs can produce an equivalent effect to a larger number of spontaneous inputs. In our model, therefore, we produce responses comparable to those produced by Wang and Colburn (2012), but with a more realistic number of inhibitory inputs.

Interestingly, our results suggest that PST discharge type does not correlate well with the characteristics observe in rate responses to SAM tones, and show only moderate correspondence with synchronization functions. These results conflict somewhat with the systematic relationship between inhibitory strength and timing, and discharge characteristics observed in model LSO responses. Our results thus do not preclude, and the weak relationship with PST discharge type may suggest an important role for, a contribution of additional mechanisms (including spontaneous inhibitory inputs) contributing to LSO unit responses, as we discussed in depth previously (Greene and Davis 2012).

### On the origin of low-pass SAM tone rMTFs

Previous results of the SFIE model (Nelson and Carney 2004, 2007), do not fully explain the tendency for units in barbiturate anesthetized cat LSO (Joris and Yin 1998) to show low-pass rMTF responses. Both the physiological and model data described above usually showed either band-pass or all-pass rMTFs. One explanation is that low- (or high-) pass curves could result from an insufficient range of modulation frequencies sampled (that is, units would appear bandpass if lower f_m_ had been recorded). This argument is sufficient to explain those results, and was necessary to explain the extraordinarily low response rates to BF tones (i.e. 0 Hz modulation) when K_LT_ is introduced to a model LSO cell (Wang and Colburn 2012). However, a second potential explanation is suggested by the asymmetry observed at some inhibitory strengths in the model rMTFs shown in Fig. 10A. Here, high inhibitory strengths (5 and above) with moderate inhibitory delays (~+2 ms) reduced responses to high f_m_ SAM tones more strongly than to low f_m_. We believe this asymmetry can potentially be explained by considering the time-course of unit responses as assessed by pure tone PST histograms: that is, by assessing the responses produced by units with an onset dominated versus a sustained dominated discharge in BF-tone PST histograms. In the following discussion, we consider the case of a unit receiving a single excitatory input with a single inhibitory input that is sufficiently strong to suppress completely output activity. In particular, we consider the effects of a strong inhibitory input with a sustained versus an onset dominated response on the output of an LSO cell receiving an excitatory input with balanced onset and sustained discharges, where both inputs arrive at the cell at the same time.

In order to explore the relative effects of the onset and sustained components of the inhibitory input, it is useful to consider the two independently. First, a cell receiving an inhibitory input with a strong sustained component will produce an output principally at the onset of the stimulus, since the responses following the initial onset response will be strongly suppressed. The unit, therefore, will produce an output once per stimulus cycle, and will produce a low discharge rate in response to low f_m_, since each cycle of the AM stimulus will appear as an independent stimulus onset. Discharge rate will thus increase with f_m_ until the stimulus period is sufficiently short that the inhibitory potential elicited by one stimulus cycle can suppress the response to the next cycle, at which point the cell’s response will be restricted to the onset (i.e. the first cycle) of the stimulus. In this manner, the cell would show a strong band-pass response. Second, when the inhibitory input is dominated by an onset component the initial rate is suppressed, but the later discharge rate may be quite high. At low f_m_, therefore, the response rate to each stimulus cycle may be high after an initial delay. With increasing f_m_ the discharge rate may increase somewhat, until the period is sufficiently short to allow the inhibition to affect subsequent cycles, at which point the cell will show strong suppression and produce essentially no response. A low-pass response, therefore, could be generated in a cell receiving an onset-dominated inhibitory input.

Inhibitory inputs to our model LSO cells are assumed to originate from LNTB, which were modeled as fast inhibitory relays of globular bushy cell projections. Furthermore, LNTB cell responses were assumed to resemble those of MNTB principal cells, since at least some LNTB cells appear to exhibit similar morphology and glycine immunoreactivity as MNTB principal cells (Spirou and Berrebi 1997). The time-course of this inhibitory input, therefore, likely closely resembles the discharge characteristics of globular bushy cells, which show primary-like with notch discharge patterns (Smith et al. 1991), and includes both a strong onset and a sustained component. When the inhibitory strength is relatively low the onset component of the inhibitory input is strong relative to the sustained component, thus the LSO cell produces a low-pass rMTF. Stronger inhibition increases the relative importance of the sustained component of the inhibition, thus shifts towards a band-pass rMTF shape. Assessing the degree to which the relative strength of onset and sustained inhibition affects rMTF shape will require future investigation particularly assessing the importance of the input integration window on LSO cell output.

### Effects of level on LSO unit MTF shape

The dependence of LSO unit responses on sound level is dramatic and consistent. Virtually all units in the current unit sample show well-tuned band-pass responses at low sound levels (near threshold), and weaker band-pass or all-pass responses at high sound levels. Similarly, synchronization was maximal at low sound levels, where the response rate was relatively low, and decreased at higher sound levels. An exception is for Onset-S units, which respond with low sustained discharge rates, and maintain high synchronization, at all sound levels. Curiously, synchronization is maintained at a relatively high value over a particular range of f_m_, even when synchronization has decreased dramatically at higher and lower f_m_, thus resulting in band-pass sMTFs at higher sound levels. These trends are recreated in the responses of the model LSO units, and, therefore, must be an effect of the relative strength and timing of excitatory and inhibitory inputs.

In most units, maximum discharge rate increases monotonically with increasing sound level to both BF-tone and BF-carrier SAM tones. Rate is typically facilitated at low sound levels for SAM tone f_m_ near the BMF, which may be due to an interaction between the periodic excitation and a post-inhibitory rebound, thereby reaching response rates above those elicited by BF-tones at the same level. With increasing sound level, however, the relative importance of this facilitation appears to decrease, and the response rates at all f_m_ increase towards the maximum discharge rate elicited by BF-tones, thus rMTF tuning is lost. Synchronization, alternatively, is maximal at low, and decreases with increasing sound level, thus well-timed responses principally appear when response rates are low. This may result because with increasing sound level, and correspondingly increasing response rates, the probability of producing an output increases independent of the AM period. Unit discharges, therefore, appear between the peaks in the SAM tone envelope, thus the vector strength correspondingly decreases. The maintenance of synchronization over a particular f_m_ range across sound levels suggests that the timing of excitatory and inhibitory inputs continues to strongly affect responses even at high levels.

### The role of LSO in the processing of AM sounds

Our assessment of LSO cell tuning to SAM tones in the decerebrate cat was initially motivated by the similarity between the feed-forward inhibition model of LSO unit discharge pattern formation (Greene and Davis 2012) and the SFIE model of AM tuning (Nelson and Carney 2004). Our results indicate that LSO units in decerebrate cat do indeed show evidence for enhanced rate tuning to AM stimuli compared to both their CN inputs, and when compared to units recorded in the LSO of barbiturate anesthetized cat. The responses, therefore, are indeed consistent with an intermediate processing stage between CN and ICC in a multi-stage SFIE type model. IN particular, while LSO units appear to show enhanced tuning compared to their bushy cell inputs (who show all-pass rate functions), response characteristics can substantial differences compared to ICC, including substantial level dependence and substantially higher rBMFs.

That LSO units show this enhanced rate tuning compared to CN, however, does not indicate whether or not the ILD processing pathway initiated by the LSO contributes towards AM processing in any meaningful way. The high fidelity coding of AM in LSO at low sound levels could merely be an epiphenomenon resulting from inhibitory mechanisms developed for processing of other stimuli (e.g. ILD); consistent with this interpretation, Koka and Tollin (2014), following from Tollin (2003), suggested that LSO must compute ILDs for biologically relevant complex stimuli, such as rapid spectrotemporally-modulated envelopes such as are present in speech/animal vocalizations, and in moving sound sources. Indeed, it is likely that alternative pathways to ICC, such as direct projections from CN, and projections through the VNLL, likely contribute a substantial or majority of the inputs responsible for AM processing. Nevertheless, LSO cells appear to show some degree of rate tuning to AM stimuli, and future work investigating responses in the ICC projection targets of the LSO (i.e. type I units (Greene et al. 2010; Ramachandran et al. 1999)) to determine whether this sensitivity is maintained within this pathway is required to further elucidate AM processing. Likewise, recordings in awake/behaving animals are necessary to determine whether descending projections (which were eliminated during the decerebration procedure) influence AM tone processing.

## ACKNOWLEDGEMENTS

We are grateful for the helpful input of Drs. L. Carney, W. O’Neill, G. Paige, and D. Tollin, as well as technical assistance from O. Lomakin and T. Bubel.

## References

Barnes-Davies M, Barker MC, Osmani F, and Forsythe ID. Kv1 currents mediate a gradient of principal neuron excitability across the tonotopic axis in the rat lateral superior olive. Eur J Neurosci 19: 325–333, 2004.

Batra R. Responses of neurons in the ventral nucleus of the lateral lemniscus to sinusoidally amplitude modulated tones. J Neurophysiol 96: 2388–2398, 2006.

Batra R, Kuwada S, and Fitzpatrick DC. Sensitivity to interaural temporal disparities of lowand high-frequency neurons in the superior olivary complex. I. Heterogeneity of responses. J Neurophysiol 78: 1222–1236, 1997a.

Batra R, Kuwada S, and Fitzpatrick DC. Sensitivity to interaural temporal disparities of lowand high-frequency neurons in the superior olivary complex. II. Coincidence detection. J Neurophysiol 78: 1237–1247, 1997b.

Boudreau JC, and Tsuchitani C. Binaural interaction in the cat superior olive S segment. J Neurophysiol 31: 442–454, 1968.

Bregman AS, Levitan R, and Liao C. Fusion of auditory components: effects of the frequency of amplitude modulation. Perception & psychophysics 47: 68–73, 1990.

Brownell WE, Manis PB, and Ritz LA. Ipsilateral inhibitory responses in the cat lateral superior olive. Brain Res 177: 189–193, 1979.

Caird D, and Klinke R. Processing of binaural stimuli by cat superior olivary complex neurons. Exp Brain Res 52: 385–399, 1983.

Cant NB, and Benson CG. Parallel auditory pathways: projection patterns of the different neuronal populations in the dorsal and ventral cochlear nuclei. Brain Res Bull 60: 457–474, 2003.

Cant NB, and Casseday JH. Projections from the anteroventral cochlear nucleus to the lateral and medial superior olivary nuclei. J Comp Neurol 247: 457–476, 1986.

Caspary DM, Backoff PM, Finlayson PG, and Palombi PS. Inhibitory inputs modulate discharge rate within frequency receptive fields of anteroventral cochlear nucleus neurons. J Neurophysiol 72: 2124–2133, 1994.

Davis KA, Hancock KE, and Delgutte B. Computational Models of Inferior Colliculus Neurons. In: Computational Models of the Auditory System edited by Meddis RSpringer, 2010, p. 129.

Dicke U, Ewert SD, Dau T, and Kollmeier B. A neural circuit transforming temporal periodicity information into a rate-based representation in the mammalian auditory system. J Acoust Soc Am 121: 310–326, 2007.

Dodla R, Svirskis G, and Rinzel J. Well-timed, brief inhibition can promote spiking: postinhibitory facilitation. J Neurophysiol 95: 2664–2677, 2006.

Evans EF, and Nelson PG. The responses of single neurones in the cochlear nucleus of the cat as a function of their location and the anaesthetic state. Exp Brain Res 17: 402–427, 1973.

Ferragamo MJ, Golding NL, and Oertel D. Synaptic inputs to stellate cells in the ventral cochlear nucleus. J Neurophysiol 79: 51–63, 1998.

Finlayson PG, and Caspary DM. Low-frequency neurons in the lateral superior olive exhibit phase-sensitive binaural inhibition. J Neurophysiol 65: 598–605, 1991.

Frisina RD. Subcortical neural coding mechanisms for auditory temporal processing. Hear Res 158: 1–27, 2001.

Glendenning KK, Baker BN, Hutson KA, and Masterton RB. Acoustic chiasm V: inhibition and excitation in the ipsilateral and contralateral projections of LSO. J Comp Neurol 319: 100–122, 1992.

Glendenning KK, Hutson KA, Nudo RJ, and Masterton RB. Acoustic chiasm II: Anatomical basis of binaurality in lateral superior olive of cat. J Comp Neurol 232: 261–285, 1985.

Glendenning KK, and Masterton RB. Acoustic chiasm: efferent projections of the lateral superior olive. J Neurosci 3: 1521–1537, 1983.

Goldberg JM, and Brown PB. Response of binaural neurons of dog superior olivary complex to dichotic tonal stimuli: some physiological mechanisms of sound localization. J Neurophysiol 32: 613–636, 1969.

Greene NT, and Davis KA. Discharge patterns in the lateral superior olive of decerebrate cats. J Neurophysiol 108: 1942–1953, 2012.

Greene NT, Lomakin O, and Davis KA. Monaural spectral processing differs between the lateral superior olive and the inferior colliculus: physiological evidence for an acoustic chiasm. Hear Res 269: 134–145, 2010.

Guinan JJ, Jr., Guinan SS, and Norris BE. Single auditory units in the superior olivary complex. I. Responses to sounds and classifications based on physiological properties. Int J Neurosci 4: 101–120, 1972a.

Guinan JJ, Jr., Norris BE, and Guinan SS. Single auditory units in the superior olivary complex. II. Locations of unit categories and tonotopic organization. Int J Neurosci 4: 147–166, 1972b.

Henning GB. Detectability of interaural delay in high-frequency complex waveforms. J Acoust Soc Am 55: 84–90, 1974a.

Henning GB. Lateralization and the binaural masking-level difference. J Acoust Soc Am 55: 1259–1262, 1974b.

Hewitt MJ, and Meddis R. A computer model of amplitude-modulation sensitivity of single units in the inferior colliculus. J Acoust Soc Am 95: 2145–2159, 1994.

Joris PX. Envelope coding in the lateral superior olive. II. Characteristic delays and comparison with responses in the medial superior olive. J Neurophysiol 76: 2137–2156, 1996.

Joris PX, Schreiner CE, and Rees A. Neural processing of amplitude-modulated sounds. Physiol Rev 84: 541–577, 2004.

Joris PX, and Yin TC. Envelope coding in the lateral superior olive. I. Sensitivity to interaural time differences. J Neurophysiol 73: 1043–1062, 1995.

Joris PX, and Yin TC. Envelope coding in the lateral superior olive. III. Comparison with afferent pathways. J Neurophysiol 79: 253–269, 1998.

Joris PX, and Yin TC. Responses to amplitude-modulated tones in the auditory nerve of the cat. J Acoust Soc Am 91: 215–232, 1992.

Koka K, and Tollin DJ. Linear coding of complex sound spectra by discharge rate in neurons of the medial nucleus of the trapezoid body (MNTB) and its inputs. Front Neural Circuits 8: 144, 2014.

Kuwada S, Batra R, and Stanford TR. Monaural and binaural response properties of neurons in the inferior colliculus of the rabbit: effects of sodium pentobarbital. J Neurophysiol 61: 269–282, 1989.

Langner G. Neuronal mechanisms for pitch analysis in the time domain. Exp Brain Res 44: 450–454, 1981.

Langner G, and Schreiner CE. Periodicity coding in the inferior colliculus of the cat. I. Neuronal mechanisms. J Neurophysiol 60: 1799–1822, 1988.

Lu H, and Xu TL. The general anesthetic pentobarbital slows desensitization and deactivation of the glycine receptor in the rat spinal dorsal horn neurons. J Biol Chem 277: 41369–41378, 2002.

MacGregor RJ. Neural and brain modeling. Academic Press, 1987.

Nelson PC, and Carney LH. Neural rate and timing cues for detection and discrimination of amplitude-modulated tones in the awake rabbit inferior colliculus. J Neurophysiol 97: 522–539, 2007.

Nelson PC, and Carney LH. A phenomenological model of peripheral and central neural responses to amplitude-modulated tones. J Acoust Soc Am 116: 2173–2186, 2004.

Plack CJ. The sense of hearing. Mahwah, N.J.: Lawrence Erlbaum Associates, 2005, p. xi, 267 p.

Ramachandran R, Davis KA, and May BJ. Single-unit responses in the inferior colliculus of decerebrate cats. I. Classification based on frequency response maps. J Neurophysiol 82: 152–163, 1999.

Rhode WS, Oertel D, and Smith PH. Physiological response properties of cells labeled intracellularly with horseradish peroxidase in cat ventral cochlear nucleus. J Comp Neurol 213: 448–463, 1983.

Rosen S. Temporal information in speech: acoustic, auditory and linguistic aspects. Philos Trans R Soc Lond B Biol Sci 336: 367–373, 1992.

Saint Marie RL, Shneiderman A, and Stanforth DA. Patterns of gamma-aminobutyric acid and glycine immunoreactivities reflect structural and functional differences of the cat lateral lemniscal nuclei. J Comp Neurol 389: 264–276, 1997.

Singh NC, and Theunissen FE. Modulation spectra of natural sounds and ethological theories of auditory processing. J Acoust Soc Am 114: 3394–3411, 2003.

Smith PH, Joris PX, Carney LH, and Yin TC. Projections of physiologically characterized globular bushy cell axons from the cochlear nucleus of the cat. J Comp Neurol 304: 387–407, 1991.

Spirou GA, and Berrebi AS. Glycine immunoreactivity in the lateral nucleus of the trapezoid body of the cat. J Comp Neurol 383: 473–488, 1997.

Steeneken HJ, and Houtgast T. A physical method for measuring speech-transmission quality. J Acoust Soc Am 67: 318–326, 1980.

Thompson AM, and Schofield BR. Afferent projections of the superior olivary complex. Microsc Res Tech 51: 330–354, 2000.

Tolbert LP, Morest DK, and Yurgelun-Todd DA. The neuronal architecture of the anteroventral cochlear nucleus of the cat in the region of the cochlear nerve root: horseradish peroxidase labelling of identified cell types. Neuroscience 7: 3031–3052, 1982.

Tollin DJ. The lateral superior olive: a functional role in sound source localization. Neuroscientist 9: 127–143, 2003.

Tollin DJ, Koka K, and Tsai JJ. Interaural level difference discrimination thresholds for single neurons in the lateral superior olive. J Neurosci 28: 4848–4860, 2008.

Tollin DJ, and Yin TC. Interaural phase and level difference sensitivity in low-frequency neurons in the lateral superior olive. J Neurosci 25: 10648–10657, 2005.

Tsai JJ, Koka K, and Tollin DJ. Varying overall sound intensity to the two ears impacts interaural level difference discrimination thresholds by single neurons in the lateral superior olive. J Neurophysiol 103: 875–886, 2010.

Tsuchitani C. Discharge patterns of cat lateral superior olivary units to ipsilateral tone-burst stimuli. J Neurophysiol 47: 479–500, 1982.

Tsuchitani C. Functional organization of lateral cell groups of cat superior olivary complex. J Neurophysiol 40: 296–318, 1977.

Tsuchitani C, and Boudreau JC. Single unit analysis of cat superior olive S segment with tonal stimuli. J Neurophysiol 29: 684–697, 1966.

Wang L, and Colburn HS. A modeling study of the responses of the lateral superior olive to ipsilateral sinusoidally amplitude-modulated tones. J Assoc Res Otolaryngol 13: 249–267, 2012.

Whitley JM, and Henkel CK. Topographical organization of the inferior collicular projection and other connections of the ventral nucleus of the lateral lemniscus in the cat. J Comp Neurol 229: 257–270, 1984.

Winer JA, Larue DT, and Pollak GD. GABA and glycine in the central auditory system of the mustache bat: structural substrates for inhibitory neuronal organization. J Comp Neurol 355: 317–353, 1995.

Wu SH, and Kelly JB. Physiological evidence for ipsilateral inhibition in the lateral superior olive: synaptic responses in mouse brain slice. Hear Res 73: 57–64, 1994.

Young ED, Robert JM, and Shofner WP. Regularity and latency of units in ventral cochlear nucleus: implications for unit classification and generation of response properties. J Neurophysiol 60: 1–29, 1988.

Zhang H, and Kelly JB. Responses of neurons in the rat’s ventral nucleus of the lateral lemniscus to amplitude-modulated tones. J Neurophysiol 96: 2905–2914, 2006.

Zhou Y, and Colburn HS. A modeling study of the effects of membrane afterhyperpolarization on spike interval statistics and on ILD encoding in the lateral superior olive. J Neurophysiol 103: 2355–2371, 2010.

Zilany MS, and Bruce IC. Modeling auditory-nerve responses for high sound pressure levels in the normal and impaired auditory periphery. J Acoust Soc Am 120: 1446–1466, 2006.

Zilany MS, Bruce IC, Nelson PC, and Carney LH. A phenomenological model of the synapse between the inner hair cell and auditory nerve: long-term adaptation with power-law dynamics. J Acoust Soc Am 126: 2390–2412, 2009.

